# CD4^+^ T_h_ immunogenicity of the *Ascaris spp*. secreted products

**DOI:** 10.1101/699231

**Authors:** Friederike Ebner, Eliot Morrison, Miriam Bertazzon, Ankur Midha, Susanne Hartmann, Christian Freund, Miguel Álvaro-Benito

## Abstract

*Ascaris spp*. is a major health problem of humans and animals alike, and understanding the immunogenicity of its antigens is required for developing urgently needed vaccines. The parasite-secreted products represent the most relevant, yet highly complex (>250 proteins) antigens of *Ascaris spp.* as defining the pathogen-host interplay. We applied an *in vitro* antigen processing system coupled to quantitative proteomics to identify potential CD4^+^ T_h_ cell epitopes in *Ascaris suum*-secreted products. This approach restricts the theoretical list of epitopes, based on affinity prediction, by a factor of ∼1200. More importantly, selection of 2 candidate peptides based on experimental evidence demonstrated the presence of epitope-reactive T cells in *Ascaris*-specific T cell lines generated from healthy human individuals. Thus, this stringent work pipeline identifies a human haplotype-specific T cell epitope of a major human pathogen. The methodology described represents an easily adaptable platform for characterization of highly complex pathogenic antigens and their MHCII-restriction.

---

*Ascaris lumbricoides*. infections currently affect around 820 million people leading to impaired growth, impaired physical fitness and cognition, while also reducing general performance, in particular in children (Jourdan *et al.*, 2018). Considering the worldwide prevalence and intensity of Ascariasis in humans, it is critical to overcome the current limitations in controlling this parasitic infection. Improvements on infrastructure and educational programs together with revised mass drug administration programs will definitively contribute to mitigate the impact of Ascariasis (Jourdan *et al.*, 2018). However, the ideal solution would be the development of vaccines that prevents commonly observed re-infection after chemotherapy (Zhan *et al.*, 2014; Noon and Aroian, 2017). Candidate vaccines to prevent *Ascaris* infection should be able to trigger effective antibody responses targeting essential antigens for the parasite to complete its life cycle (Harris and Gause, 2011). The challenge is, that these large, multicellular and cuticularized parasites confront the host with a complex mixture of protein antigens from which we have still very poor knowledge on their importance for infection or their immunogenicity. *Ascaris* parasites actively excrete and secrete complex mixtures of molecules, *Vic*. the Excretory Secretory (ES) products, which are essential in parasite’s communication with its host, and shaping the host immune response (Hewitson, Grainger and Maizels, 2009; Wang *et al.*, 2013; Chehayeb *et al.*, 2014). The ES proteins comprise critical targets for vaccination in animal models and are expected to bear targets for vaccination in humans as well (reviewed in (Zhan *et al.*, 2014)). Intriguingly, a considerable heterogeneity was observed in antibody profiles of human infected individuals (Haswell-Elkins *et al.*, 1989; Kennedy *et al.*, 1990), and work on animal models demonstrated the Major Histocompatibility Complex of class II-control of *Ascaris* antibody repertoires (Tomlinson *et al.*, 1989; Kennedy, Fraser and Christie, 1991). Thus, the deterministic nature of T-B cell responses - which implies that antigen-specific B cells receive help from antigen-specific, activated CD4^+^ T_h_ cells - should contribute to define targets for the design of candidate subunit vaccines. Therefore, identifying the epitopes that are involved in the host’s natural CD4^+^ T_h_ cell responses is essential to understand, monitor or modulate adaptive immune mechanisms that orchestrate *Ascaris spp*. expulsion. However, the high complexity of *Ascaris* ES products and the polymorphic nature of Major Histocompatibility Complex of class II (MHCII) molecules represent two major challenges to define immunogenic antigens for a majority of humans.

CD4^+^ T_h_ cells recognize antigenic peptides presented by the major histocompatibility complex class II (MHCII) proteins expressed on antigen-presenting cells (APC). Peptides from antigens only become immunogenic when they are selected for presentation, and remain bound to MHCII molecules for a sufficient time to allow T cell surveillance. Thus, the abundance of antigens, their resistance to degradation and the affinity for the MHCII will define the potential immunogenicity of any peptide. To date, conventional *in silico* approaches predict peptide-MHCII affinity mostly based on MHC pocket occupation by the peptide amino acids (Lundegaard, Lund and Nielsen, 2012; Sanchez-Trincado, Gomez-Perosanz and Reche, 2017). However, these approaches ignore relevant aspects affecting epitope selection, such as the dynamics of peptide-MHCIIs (Wieczorek *et al.*, 2016, 2017) (pMHCII), the peptide-editing function of HLA-DM (Álvaro-Benito *et al.*, 2014), and the influence of proteolytic activities on antigen presentation (Kim and Sadegh-Nasseri, 2015). Although integrating proteolytic degradation improves the current conventional methods (Barra *et al.*, 2018), a robust *in silico* ranking of the most effectively presented peptides is still elusive. Experimental approaches based on recombinant proteins or subcellular fractions containing endosomal compartments rich on MHCII have been applied to define epitope selection on single antigens (Hartman *et al.*, 2010; Mutschlechner *et al.*, 2010). Culture of primary DC and quantitative immunopeptidomics of infected cells has also been used to define CD4^+^ T_h_ cell epitopes from *Listeria monocytogenes* in mouse (Graham *et al.*, 2018). However, to date, there is no experimental set up to define the CD4^+^ T_h_ immunogenicity of complex antigenic mixtures in humans.

Here, we introduce a bottom-up experimental approach to delineate the CD4^+^ T_h_ immunogenicity of *Ascaris* ES. As antigen source we used ES from *Ascaris suum*, the closely related porcine analogue of *A. lumbricoides*, which is considered antigenically identical. Initially, we performed a proteomic characterization of male and female parasite ES (ESM and ESF respectively) products (Wang *et al.*, 2013; Chehayeb *et al.*, 2014) to define the antigens bearing potential CD4^+^ T_h_ epitopes (Fig.1A). By combining the use of the exponentially modified Protein Abundance Index (emPAI) with ^16^O/^18^O-labelling we defined the ESM and ESF composition, thereby retrieving absolute protein abundances of ESM and ESF extracts (Fig. 1B left), and the relative difference between them (Fig. 1B right). This analysis yielded additional information regarding the differences in abundance between ESF and ESM (Table S1) in comparison to the previously described dataset (Chehayeb *et al.*, 2014). The difference in the number of protein sources found here when compared with the previous study (175 previously *vs.* 254 in our case) arises from our conservative approach when considering the source of tryptic peptides. Rather than selecting a single leading protein, we explicitly kept protein entries with small differences in their primary sequences but which are not clearly distinguishable by conventional shotgun MS. This characterization provides a detailed overview of the potential antigenic sources of CD4^+^ T_h_ cell epitopes. We found large differences in the abundance of proteins either within a single extract or between ESM and ESF (fold-differences up to 10^3^ for emPAI and 10^7^, respectively). We annotated all entries according to the defined absolute and relative abundance to further evaluate whether protein abundance is a key factor defining the immunogenicity of ES proteins.

**Fig. 1.**
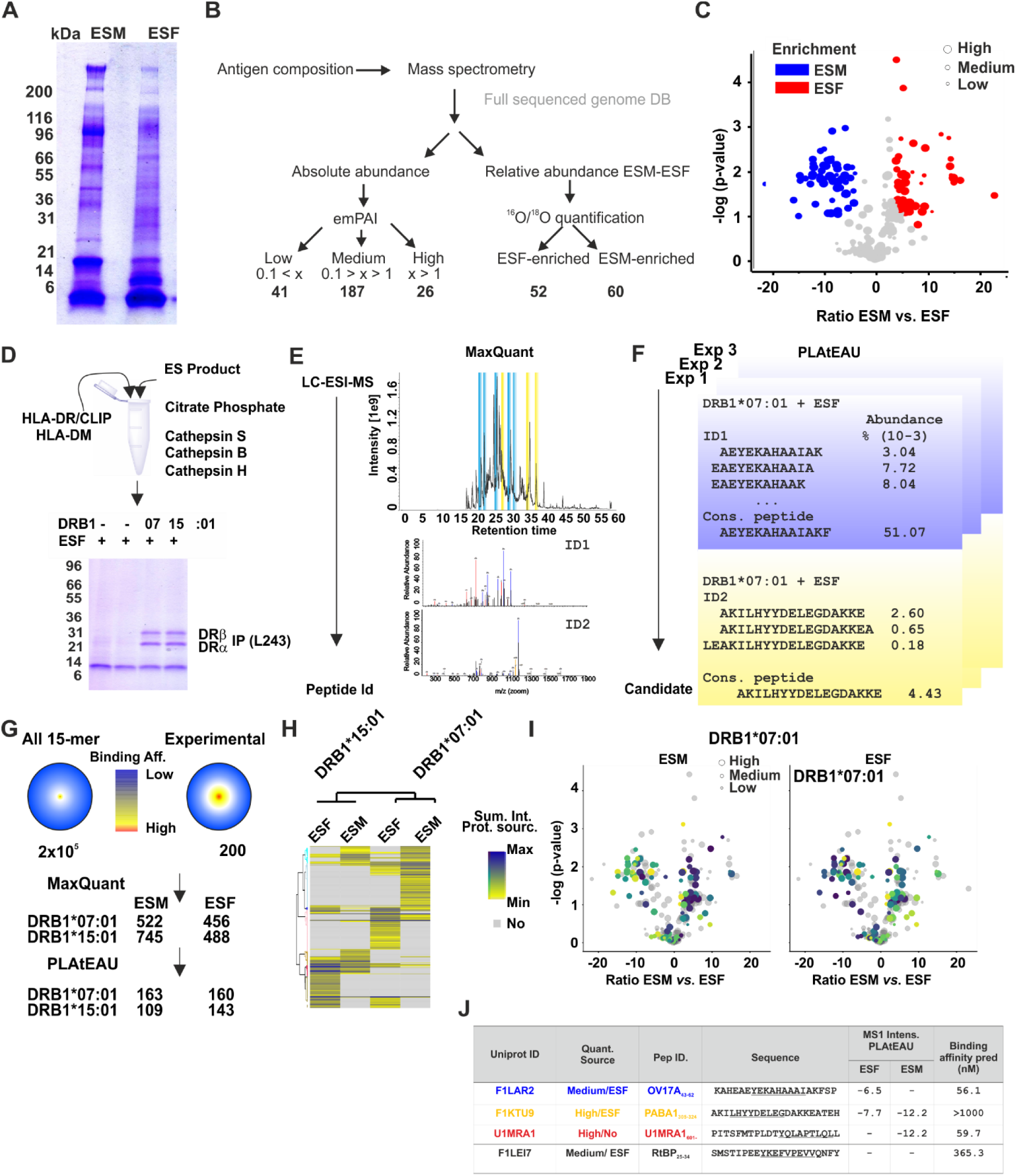
Assessing potential CD4^+^ T cell epitopes from complex antigenic mixtures. **A**. Model antigenic mixtures derived from *Ascaris* ES products are different in protein composition. SDS-PAGE of ES male (ESM) and ES female (ESF) mixtures (40 μg of antigen loaded per well). **B.** A mass-spectrometry-based approach used to determine composition of ES products. The emPAI and the ESF *vs*. ESM (^16^O/^18^O) ratios are defined after tryptic digest defining the total abundance and the differences between ESF and ESM. **C**. Volcano plot showing differences in protein composition determined by mass spectrometric analysis. The depicted log2-fold intensity difference (ESM *vs.* ESF) represents the difference in geometric mean values from ^18^O labelling. On the y axis the –log(p-value) for the difference observed is indicated. Blue symbols indicate ESM-enriched and red corresponds to ESF-enriched. The different size of the corresponding symbols indicates the determined emPAI value as shown in the legend. **D**. ES products are incubated *in vitro* with recombinant proteins and adequate buffer conditions. Proteolytic activities used degrade most of the components of the reactions except for MHCII proteins which are subsequently pull down using a conformation specific antibody (L243) **E.** MaxQuant is used for their identification. **F**. PLAtEAU is used to define series of nested peptides and retrieve the consensus peptides and corresponding MS1 intensities according to the MaxQuant output. For each peptide a relative abundance value is retrieved based on the MS1 intensity and the total ion current from the run. This approach yields a list of candidate epitopes with relative abundance values and predicted affinities. **G.** The performance of the approach is graphically represented as the average of the two allotypes and two distinct ES extracts used. ES extracts bear a total of 2 × 10^5^ potential 15-mers with a distribution of affinities shown as indicated by the legend. The data reduction of our experimental and data processing scheme is shown. **H.** Heatmap after clustering analysis of the potential candidates selected for each condition after removing background binders. The relative abundance of each candidate is shown according to the legend. **I.** Mapping of the identified potential epitopes to their corresponding protein sources. Colored dots (according to legend in E) indicate the protein sources that yield experimental epitopes. **J.** A defined set of 4 potential epitopes were selected to evaluate their potential immunogenicity. The summed intensity MS1 (as log2) is indicated for each condition as well as the predicted binding affinity for the corresponding HLA allotype. When no value is indicated, the potential epitope was not found.

We then hypothesized that the potential CD4^+^ T_h_ epitopes of *Ascaris* secreted products could be experimentally defined by combining key events of HLA-specific antigen presentation as described by the minimalist approach of Sadegh-Nasseri’s group (Hartman *et al.*, 2010). Using this system, the lower background of self-peptides will benefit the MS identification of pathogen derived antigenic peptides. We applied the ESM or ESF antigens, two common MHCII allotypes in *Ascaris*-endemic territories (DRB1*07:01 or DRB1*15:01, see Fig. S1A) preloaded with the placeholder Class II invariant chain peptide-CLIP and HLA-DM, which functions as peptide-editor (Álvaro-Benito *et al.*, 2014) (details in Fig. S1B) and the commercial proteases previously described (Hartman *et al.*, 2010). To evaluate the impact of protein composition on epitope selection the distinct antigen profiles of ESF and ESM were used. Control experiments confirmed that ES antigens are degraded by cathepsins while MHCII are resistant (Fig. 1D). Immunoprecipitation of HLA-DR molecules and treatment with TFA yielded the pool of bound peptides for each condition, which were further analyzed by LC-ESI-MS. We used MaxQuant for peptide identification considering a customized database that includes all the entries found in the ES products and the molecules of the *in vitro* reconstitution system (MHCII, HLA-DM and proteases). The information on MS1 intensity for each identified peptide (Fig. 1E) was used by our recently described epitope analysis tool, PLAtEAU (Álvaro-Benito *et al.*, 2018), to define consensus peptides including their relative abundance for each series of peptides that contain a common core but vary in the length of their N- and C-terminal extensions (Fig. 1F). The full list of 335 potential of CD4^+^ T cell epitopes was annotated based on the abundance of the corresponding protein source and the predicted affinity for each allotype (Table S2).

Our experimental workflow leads to a massive reduction in the number of potential epitopes from a total of 2 × 10^5^ potential 15-mer peptides in the whole ES protein extract, to less than 10^3^ peptides, which once analyzed with PLAtEAU resulted in around 150 candidate epitopes for each condition (details given in Fig. S1B). Furthermore, the presence of predicted weak and high affinity binders increases from 10% and 1 % in the pool of all potential peptides of the ES to 20 % and 7% in the experimentally determined peptides, respectively (details given in Fig. S1B). Surely, the inclusion of HLA-DM in the system is a major driver for the focusing of the repertoire as it is seen here (Fig. 1G). Hierarchical clustering analysis of the annotated potential epitopes based on their abundance reveals that both MHCII molecules selected mostly epitopes from proteins with intermediate and high emPAI values, and in particular from those enriched (e.g. ESF:ESM ratio > 2) in the respective antigen source (ESM or ESF) (Fig. 1H). Mapping the intensities to their sources in the volcano plots depicting the compositional overview of the ES confirmed the result (Fig. 1I and S1C). Most interestingly, we found the Ov17_43-64_ consensus peptide as a prime candidate to test its potential immunogenicity for two main reasons. First, this peptide it is selected exclusively under DRB1*07:01 + ESF conditions, thereby representing an ideal candidate to prove the selectivity and performance of our experimental approach. Second, experiments on swine and mouse models have shown the potential of Ov17 antigen (also referred to as As16) for conferring protection to *Ascaris suum* infection (Tsuji *et al.*, 2003, 2004), and for eliciting Th2-type responses (Wei *et al.*, 2017) indicating its interpecies immunogenicity. Furthermore, in humans, the restricting DRB1*07:01 allele reaches up to 9 % of the global population (Gourraud *et al.*, 2014) and even higher frequencies on *Ascaris lumbricoides* endemic areas with up to 15 % in specific African and Asian populations.

We were interested in validating whether the Ov17_43-67_ candidate is recognized as a human T cell epitope, and evaluating the performance of the reconstituted *in vitro* system, by applying the TcR pool present in healthy individuals to a limited set of candidates. Our restricted set of candidates includes one peptide with a similar affinity for DRB1*07:01 but not selected from the ESF antigen (U1MRA1_601-620_), a peptide (PABA1_305-324_) that belongs to the immunogenic Polyprotein ABA1 (Kennedy, Fraser and Christie, 1991; Kennedy *et al.*, 2008), and a peptide (RtBP_25-45_) not selected by this allotype. Thus, out of the four peptides chosen (Fig. 1J), only OV17_43-62_, should be selected for presentation from ESF antigen, and therefore elicit T cell responses when presented by DRB1*07:01. We generated human T cell lines from healthy volunteers reactive against the whole ESF antigen using antigen-specific T cell enrichment and expansion as described by Bacher *et al*. (Bacher *et al.*, 2013) (Fig. 2A). This approach is based on the up-regulation of CD40-L shortly after TcR-mediated antigen recognition irrespectively of the restricting MHC allele and can be used to assess and enrich antigen-specific T cells (Bacher and Scheffold, 2015). In our case, it helped us to overcome the expected low *in vivo* frequency of any potential *Ascaris* ESF-specific CD4^+^ T_h_ cells in healthy (uninfected) donors, as confirmed by CD40-L staining following ES antigenic stimulation (Fig. 2B). Indeed, a considerable amount of CD40-L^+^ cells (mean ± SEM: 33.8 % ± 5.3 among CD4^+^ T cells) resulted from re-stimulation of enriched and expanded human cells with *Ascaris* ESF antigen loaded to APCs compared to unloaded autologous APCs (Fig. 2C and 2D). This refers to the population of total ES-reactive CD4^+^ T_h_ cells of different donors after expansion. Upregulation of CD40-L and CD40-L/cytokine co-expression after re-stimulation confirms a functional CD4^+^ T_h_ phenotype of the generated cell lines (Fig. S2).

**Fig. 2.**
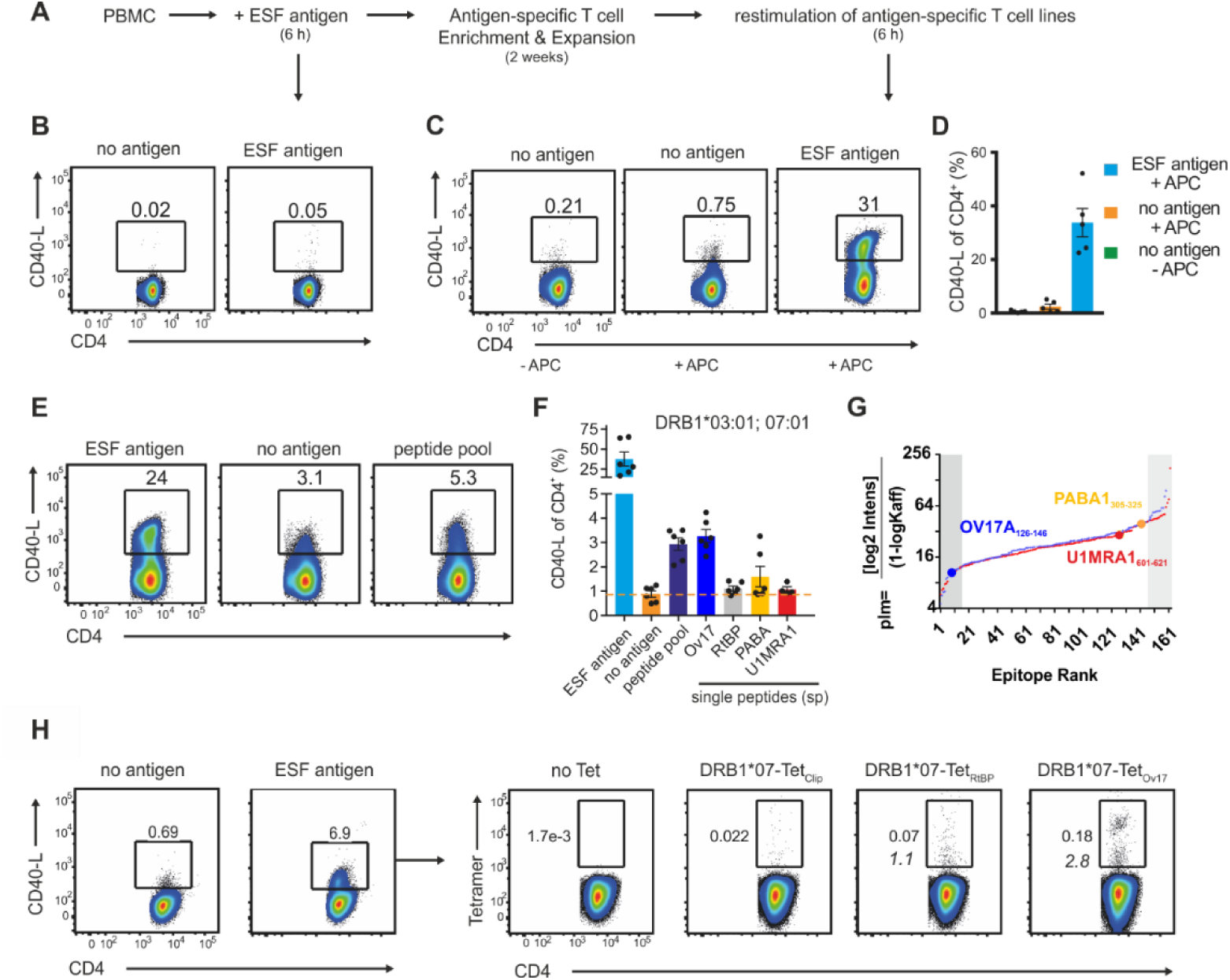
HLA-selected epitopes induce CD4^+^ T cell activation in *Ascaris* ES antigen-specific T cell lines. **A.** Experimental overview of generating *Ascaris* ES-specific T cell lines. PBMCs from healthy donors were stimulated with 40µg/ml ES antigen F for 6h, enriched for CD154^+^ cells (CD40-L^+^) and expanded for 2 weeks. **B.** Frequencies of CD40-L expressing cells among CD4^+^, representing *Ascaris* ES-specific T cells, were analyzed prior to expansion. **C.** Same as in B after re-stimulation with or without ESF antigen-primed, CD3-depleted APC and are indicated by the numbers above gates. **D.** Indicated are percentages of CD40-L/CD154^+^ antigen-reactive T cells among CD4^+^ cells after expansion and re-stimulation with ESF antigen for n=5 different donors. **E.** Representative dot blot example for a ESF antigen-specific T cell line generated from an healthy DRB1*03:01;07:01-typed volunteer and re-stimulated with either whole ES antigen (40 µg/ml), no antigen or a pool of synthetic peptides (25 µg/ml for each peptide) selected from the *in vitro* reconstituted HLA-DRB1*07:01 experimental data set (Fig. 2F). **F.** Summarizes CD40-L frequencies among CD4^+^, indicative for peptide recognition by CD4^+^ T cells, for whole ES antigen, peptide pool and single peptide (sp) re-stimulations. Combined are data from the same healthy, DRB1*03:01;07:01-typed volunteer from n=3 separate experiments with n=2 separate re-stimulations. CD40-L frequencies per experiment were corrected for individual background CD40-L expression of no antigen/ - APC controls. **G.** Based on the experimental data from Fig 2. We ranked the potential epitopes defined by our experimental system according to the coefficient between total abundance of the corresponding epitope, and its predicted affinity. The obtained value was define as predicted immunogenicity. Graphical representation of pIm values for all epitopes identified for each ES antigen (ESM in blue and ESF in red) using DRB1*07:01. Peptides indicated in different colors and larger datapoints correspond to those selected for T cell experiments. Shaded areas represent the highest and lowest deciles of pIm values. **H.** Representative dot blots of an *Ascaris* ESF-specific, DRB1*03:01;07:01 T cell line analyzed for *Ascaris* ESF peptide-specific tetramer staining. Left side indicates overall frequency of ESF antigen specific CD4^+^ cells after expansion compared to control. Right side shows corresponding tetramer staining with DRB1*07-Tet-_CLIP_ (control), Tet-_RtBP_ and Tet_Ov17_ gated on CD4^+^ T cells after expansion. Italic numbers indicate calculated Tet^+^ frequency relative to proportion of ESF antigen-specific T cells.

Furthermore, we derived ESF T cell lines from a healthy, DRB1*03:01;07:01-typed volunteer and assessed CD40-L expression after re-stimulation with either whole antigen, the selected pool of four peptides, or single peptide loaded APCs (Fig. 2E). Strikingly, we found that the Ov17_43-62_, the only candidate selected as potential CD4^+^ T_h_ cell epitope by our approach with predicted high affinity, yield a stronger T cell response than any of the other peptides assayed (Fig. 2F). These results therefore confirm the predictive power of the reconstituted *in vitro* system. Indeed, the combination of all peptides as well as only the single peptide Ov17_43-62_ generated a CD40-L response above the background (shown as a dashed line).

We reasoned that we could derive a parameter to define the potential immunogenicity (pIm) of all experimentally defined epitopes based on their measured abundance. This parameter reflects the ratio between the absolute value of the log2 MS1 intensity for the epitope (lower for high abundant) and the 1-logKaff value as its predicted affinity (increases with higher affinity). Thus, the lower the pIm value for a candidate, the higher the likelihood to be immunogenic as indeed is reflected for the pool of potential epitopes selected (Fig. 2G). Indeed, for the two candidates found in DRB1*07:01 + ESF, the pIm value reflects the abundance of T cells reactive after expansion with the whole ESF.

Considering the co-dominant expression of other HLA genes in the donor-derived cell lines, the reactivity observed could arise from the presentation of the corresponding peptides by any other donor-MHCII allotype. We confirmed the DRB1*07:01 restriction for the presentation of the Ov17 peptide by tetramer staining of ESF CD4^+^ T cell lines. A significant pool of CD4^+^ T cells responding to this peptide is detected when it is displayed by DRB1*07:01 tetramers compared to control tetramers (DR7_CLIP_ or DR7_RtBP25-44_ tetramers; Fig. 2H). Of note, the overall low frequency of DRB1*07:01-Tet_Ov17+_ cells can be explained by the low frequency of whole antigen-reactive T cells for the cell line applied in that assay (only 6.9% CD40-L^+^, Fig. 2G), but still reflect that 2.8% of all ESF reactive T cells bind DRB1*07:01-Tet_Ov17_.

Together, we demonstrate that a reduced set of *in vitro* and *ex vivo* experiments is extremely useful to define human CD4^+^ T_h_ cell epitopes from complex antigenic mixtures bypassing the need of animal models (Walsh *et al.*, 2017) or immunization in humans (Gilchuk *et al.*, 2013). Furthermore, our system represents an excellent platform for testing the importance of specific antigen processing factors such as HLA-DM (Alvaro-Benito *et al.*, 2018) or its competitive inhibitor HLA-DO (Denzin *et al.*, 2017). Conditions can be tuned to include distinct MHC allotypes or combinations thereof, and the nature of the antigen can reach from single proteins to complex mixtures derived from secretomes, complete pathogens or cellular lysates. From a biological point of view we characterize the Ov17_43-62_ epitope as a human CD4^+^ T_h_ cell epitope for *Ascaris* and show its restriction by the DRB1*07:01 allotype. Immunization with this antigen in animal trials elicits considerable protective immunity to *Ascaris suum* including specific antibodies and Th2-type cytokine responses (Tsuji *et al.*, 2003, 2004; Wei *et al.*, 2017). It will thus be of great importance to further characterize immune responses against the Ov17/As16 antigen in other MHCII allotypes to define its potential for vaccination. Another interesting candidate for instance is PABA1 (Kennedy *et al.*, 2008), showing a completely different peptide selection pattern for the two alleles used, with a considerably higher number of consensus peptides selected by DRB1*15:01 when compared to DRB1*07:01 (7 *vs.* 1 respectively). Prospectively, the stage is then set to investigate a pooled, yet limited number of antigens to profile the T cell immune status of infected individuals.. Gaining access to HLA-typed material will contribute to define the deterministic nature of B-T cell responses to complex antigens to rationalize the development of vaccine subunits to *A. lumbricoides* and *A. suum*.

## Supporting information

Table S1. Identified proteins

Table S2. Consensus peptides

## Author Contributions

M.A-B. and F.E. conceived the work, perfomed experiments, analyzed data and wrote the manuscript. E.M. performed the MS measurements and analyzed MS data, M.B. perfomed *in vitro* reconsitutued MHCII antigen processing system experiments, A.M. contributed to T cell experiments, S.H., and C.F. contributed with materials, infrastructure, scientific discussions and helped to shape the final manuscript.

## Acknowledgments

For mass spectrometry, we would like to acknowledge the assistance of the Core Facility BioSupraMol supported by the Deutsche Forschungsgemeinschaft (DFG). C.F. is thankful for funding by the DFG (FR-1325/17-1, TRR-186). The study was further supported by the DFG funded national research training group (GRK) 2046 to SH.

## Materials and methods

### Antigen preparation

Excretory-secretory (ES) antigens were prepared from worm culture supernatants of male and female adult *Ascaris suum* worms obtained from a local slaughter house. In brief, worms were separated by sex and washed several times in a balanced salt solution (BSS) containing antibiotics and used as culture media for adult worms (127 mM NaCl, 7.5 mM NaHCO_3_, 5 mM KCl, 1 mM CaCl_2_, 1 mM MgCl_2_, 200 U/mL penicillin, 200 μg/mL streptomycin, 50 μg/mL gentamicin, 2.5 μg/mL amphotericin B) and kept at 37°C with 5% CO_2_. Media was replaced on a daily basis, sterile filtered through a 0.22 μM vacuum-driven filter system and collected for ES antigen preparations starting 48h after beginning of worm culture and finally stored at −20°C until further use. Worm culture supernatants collected over 1 week were further concentrated using centrifugal protein concentrators with a 5 kDa MWCO (Vivaspin, Sartorius, Göttingen, Germany) to obtain the final, concentrated ESF antigen (from female worms) and ESM (from male worms).

### Mass spectrometry

Peptide mixtures were analyzed by a reversed-phase capillary system (Ultimate 3000 nanoLC) connected to an Orbitrap Velos (Thermo Fischer) using conditions and settings described previously(Álvaro-Benito *et al.*, 2018).

For quantitative proteomics, we used ^16^O/^18^O-labeled samples. Male and Female ES products (ESM and ESF, 50 μg) were resolved in SDS-PAGE (4-20%). Each lane was cut into ten pieces of equal size in a parallel fashion. In gel tryptic digestion and ^16^O/^18^O labelling was performed as previously described (Morrison 2015). Briefly, gel bands were incubated with 50 ng trypsin in 15 μl 50 mM ammonium bicarbonate buffer in the presence of heavy (H_2_^18^O) and light (H_2_^16^O) water overnight at 37 °C. Residual trypsin activity was inactivated by adding 10 μl of 0.5% TFA in acetonitrile. Matching heavy and light samples from either Male and Female ES were mixed before drying the samples in a Speed-Vac. Samples were reconstituted in 10 μl 0.1% (v/v) TFA and 5% (v/v) acetonitrile before measurements. Peptide identification was performed using Mascot Distiller (version 2.4.3.3) software. Data were searched against the Uniprot *Ascaris suum*. protein database (March 2017). The mass tolerance of precursor and sequence ions was set to 10 ppm and 0.35 Da, respectively. For quantification, only unique peptides were used.

Peptides eluted from the reconstituted *in vitro* antigen processing system were measured as described and MaxQuant software (version 1.5.2.8) was used for peptide identification. Customized databases featuring reviewed and non-redundant Uniprot *Ascaris* spp. proteins from uniprot were used (accessed March 2017) for the *in vitro* reconstituted experiments to which we included all other recombinant proteins used in the assay, namely human cathepsins, HLA-DR2 and -DR7, as well as HLA-DM. No enzyme specificity was used for the search, and a tolerance of 10 ppm was allowed for the main ion search and 50 ppm for the MSMS identification The “match between runs” feature was enabled. The FDR was set at 0.01 (1 %). Reverse IDs and known contaminants like keratins were filtered before further data analysis. Raw files and processed data is available upon request and will be deposited in the PRIDE repository.

### Bioinformatics prediction of MHCII binding

NetMHCIIPan 3.2 was used to predict peptide binding to the indicated HLA-DR allotypes. The protein database generated upon the MS analysis (only quantified proteins) was loaded into the webserver using a 2 percent and 10 percent cutoff for strong and weak binders.

### Constructs, protein expression and purification

DNA constructs encoding HLA molecules used in this study have been generated according to the sequences available in the IMGT/HLA database (http://www.ebi.ac.uk/ipd/imgt/hla/). The cDNAs encoding the different HLA subunits were cloned into the pFastBacDual. Recombinant proteins were expressed in Sf9 cells infected at an MOI of 5 for 72 h. Supernatants were treated as previously reported to purify the target proteins by immunoaffinity chromatography using anti-HLA-DR-FF-sepharose or M2 anti flag (Sigma)(Álvaro-Benito *et al.*, 2014).

Depending on the application, HLA-DR molecules were further treated prior to their use with only Thrombin (20 U/mg protein; for tetramer preparation) or with Thrombin and V8 protease (10 U/mg; *in vitro* reconstituted system) for 2 h at 37 °C. Subsequently proteases were inactivated by adding Complete protease inhibitor cocktail (Roche) and further gel-filtrated, while HLA-DM proteins were directly subjected to gel filtration. Both types of HLA proteins were gel-filtrated in a Sephadex S200. Fractions containing the proteins of interest were pooled and concentrated with Vivaspin 10 KDa MWCO spin filter.

### Peptide selection and synthesis

The complete list of peptides obtained after PLAtEAU analysis for all experiments was loaded into excel as a single list. We used the excel random selection function to generate a list of 4 peptides from the whole dataset. The corresponding sets of 4 peptides (500 iterations) were screened to define those with two peptides originating from the same antigen (ESF or ESM) and found in one set of experiments (DRB*01:07 or DRB*15:01). An additional criteria included the presence on the set of 4 peptides of at least one peptide not found in the corresponding experiment (used as control). The corresponding list of peptides indicated arises as the one fulfilling these criteria and having the larger distance between pIm values. Synthetic peptides were subsequently purchased from Peptides and Elephants (Berlin, Germany). Purity as stated by the vendor was more than 95 %. All peptides were protected in their N- and C-termini by addition of a Ac and NH_2_ group respectively.

### *In vitro* reconstituted antigen processing system

The previously described cell-free reconstituted *in* vitro system(Hartman *et al.*, 2010) was modified according to the specific needs of the experiments. HLA molecules together with the candidate antigens and HLA-DM were incubated for 2 h at 37° C in citrate phosphate 50 mM pH 5.2 in the presence of 150 mM NaCl. Cathepsins were added to reaction mixtures after incubation with L-Cysteine (6 mM) and EDTA (5 mM). Cathepsin B (Enzo), H (Enzo), L (Enzo) and S (Sigma) were used for our *in vitro* experiments at molar ratios (cathepsin:substrate) ranging from 1:250 to 1:500. The final reaction mixture was incubated at 37° C for 2 to 5 hours. Afterwards the pH was adjusted to 7.5 and Iodoacetamide was added (25 μM). Immunoprecipitation (IP) of the pMHCII complexes was performed using L243 covalently linked to Fast Flow sepharose. Peptides were eluted from purified MHCII adding TFA 0.1% to the samples. Peptides were separated from the MHCII molecules by using Vivaspin filters (10kDa MWCO) and a subsequent reverse phase C18 enrichment. The filtrates were further lyophilized and resuspended for mass spectrometry analyses in a mixture of H_2_O (0.94):AcN (0.05): TFA (0.01).

### Cell isolation and stimulation

Permission for experiments with human primary cells with the consent of healthy donors was obtained from the local ethic committee. Peripheral blood mononuclear cells (PBMC) were isolated by density gradient centrifugation over human Pancoll (1.077 g/ml, PAN-Biotech, Aidenbach, Germany) and rested over-night in serum-free RPMI-1640 supplemented with 100 U/ml penicillin and 100 µg/ml streptomycin (all from PAN-Biotech, Aidenbach, Germany). For enrichment, 2 × 10^7^ PBMC/ml/well were stimulated for 6h with excretory-secretory antigen mixtures of female *Ascaris suum* worms (ESF, 40 µg/ml) in the presence of anti-CD40 (clone HB14, 1 µg/ml functional grade pur, Miltenyi Biotec, Bergisch Gladbach, Germany) and anti-CD28 (clone CD28.2, 1 µg/ml, BD Pharmingen™, NJ, US). Following stimulation, antigen-specific T cells were separated using the CD154 MicroBead Kit (Miltenyi Biotec, Bergisch Gladbach, Germany) according to the manufacturer’s suggestions. That included labeling first with CD154-biotin and magnetically anti-biotin microbeads in a second step followed by enrichment using MS MACS columns (Miltenyi Biotec, Bergisch Gladbach, Germany).

For assessing precursor frequencies of *Ascaris* ESF specific CD4^+^ T cells, PBMC were stimulated as described above and Brefeldin A (3 µg/ml, Thermo Fisher Scientific) was added after the first 2h of stimulation before intracellular staining for CD40-L.

### Generation of *Ascaris suum* ES antigen-specific T cell lines and restimulation

The generation of Ag-specific T cell lines was performed as previously described(Bacher *et al.*, 2013). In brief, the stimulated and isolated CD154^+^ *Ascaris* ES antigen F-specific T cells were cultured 1:100 with autologous, mitomycin C (Sigma-Aldrich) treated feeder cells in X-VIVO™ 15 (Lonza, Basel, Switzlerland) supplemented with 5% human AB serum (PAN-Biotech, Aidenbach, Germany), 100 U/ml penicillin, 100 μg/ml streptomycin (PAN-Biotech, Aidenbach, Germany), and 50 ng/ml IL-2 (PeproTech, NJ, US). Cells were expanded for 14 days and culture medium was replenished with IL-2 containing media when needed. For restimulation after 14 days, autologous PBMC were CD3-depleted using BD FACSAria™ III cell sorter and co-cultured 1:1 with expanded T cell lines in the presence of the indicated antigens. For assessing the frequency of total *Ascaris* ESF reactive T cells after expansion, co-cultured cells were restimulated with 40 µg/ml ESF for 6h. For addressing peptide specificities, restimulation was performed with single, synthetic peptides (25 µg/ml) or a pool of all peptides (25 µg/ml of each peptide). Brefeldin A (3 µg/ml) was added after the first 2h of stimulation.

### Antibody staining and flow cytometric analysis

The following monoclonal antibodies reactive with human species were used for staining: CD8-VioGreen (BW135/80, cat.: 130-113-726), CD4-APC-Vio770 (VIT4, cat.: 130-113-211), CD154/CD40L-PE-Vio770 (5C8, cat.: 130-096-793), CD20-VioGreen (LT20, cat.: 130-113-379), CD14-VioGreen (TÜK4, 130-113-153) (all from Miltenyi Biotec, Bergisch Gladbach, Germany);

IL-13-FITC (PVM13-1, cat.: 11-7139-42 or 85BRD, cat.: 11-7136-42), IL-4-PE (8D4-8, cat.: 12-7049-42) (all from Thermo Fisher Scientific); TNFa-Pacific Blue (MAb11, cat.: 502920), IFNg-PerCp-Cy5.5 (4S.B3, cat.: 502526) (all from Biolegend, San Diego, CA, US); CD3-APC (Okt3, cat.: 17-0037-42, BD Biosciences). Fixation and permeabilization were performed using the Foxp3 Transcription Factor Staining Buffer Set (Thermo Fisher Scientific) according to the manufacturer’s suggestions followed by intracellular cytokine and CD154/CD40-L staining. Fixable viability dye was used in eFluor-506 (cat.: 65-0866-14, Thermo Fisher Scientific). Cells were acquired using BD FACSCanto II (with Diva software, Heidelberg, Germany) and post-acquisition data analysis was carried out using FlowJo software (TreeStar, Ashland, OR, US).

### Tetramer preparation and staining

Purified recombinantly expressed HLA molecules were treated with Thrombin and subsequently subjected to size exclusion chromatography (Sephadex S200). The placeholder peptide CLIP was exchanged by the indicated peptides incubating HLA molecules with 50 molar excess of the desired peptide for 72 h in the presence of molecular loading enhancers. After gel filtration the peptide loading of HLA-II complexes was verified by MS. The generated peptide HLA class II complexes were biotinylated in a BirA sequence (DRB chain) using a BirA ligase (Avidity). The Biot-peptide-HLA class II complexes were then used to generate tetramers using Streptavidin-PE. Tetrameric complexes were finally separated by gel filtration and stored in PBS + NaAz (0.02%).

For peptide-specific tetramer staining, expanded cells were incubated in sodium azide (5mM) containing X-VIVO™ 15 (Lonza) supplemented with 5% human AB serum (PAN-Biotech, Aidenbach, Germany), 100 U/ml penicillin, 100 μg/ml streptomycin (PAN-Biotech, Aidenbach, Germany) for 2h at 37°C and 5% CO_2_ in the presence of the PE-SAv-Tetramers specific for CLIP, Ov17 or RtBp at final concentrations of 20 µg/ml. Following incubation, cells were washed and counterstained for CD8, CD4, CD20, CD14 and viability.

### Statistical analysis

GraphPad Prism 7.0 software (GraphPad Software San Diego CA, USA) was used in general for statistical analysis. Variance was calculated with the 2-way ANOVA method. The null hypothesis was rejected when the p value was lower than 0.05.

Perseus software(Tyanova *et al.*, 2016) was mainly used to analyze the MS data. Epitopes identified by the PLAtEAU algorithm (% Intensity from the TIC) were loaded as matrixes into Perseus. Data was log2 transformed and missing values were imputed as 0. The resulting matrices were plotted as heat-maps. Columns were hierarchical clustered with “average” as agglomeration method and “Pearson correlation” as distance matrix. Rows were ordered by hierarchical clustering using “average” as agglomeration method and “euclidean” as distance matrix. Epitopes eluted from each experimental condition were grouped and used to define the mean intensity value for each peptide or epitope. P values were calculated based on the observed intensities using t-test, in this case an FDR of 0.01 and a S0 = of 2 were used.

---

**Fig. S1.**
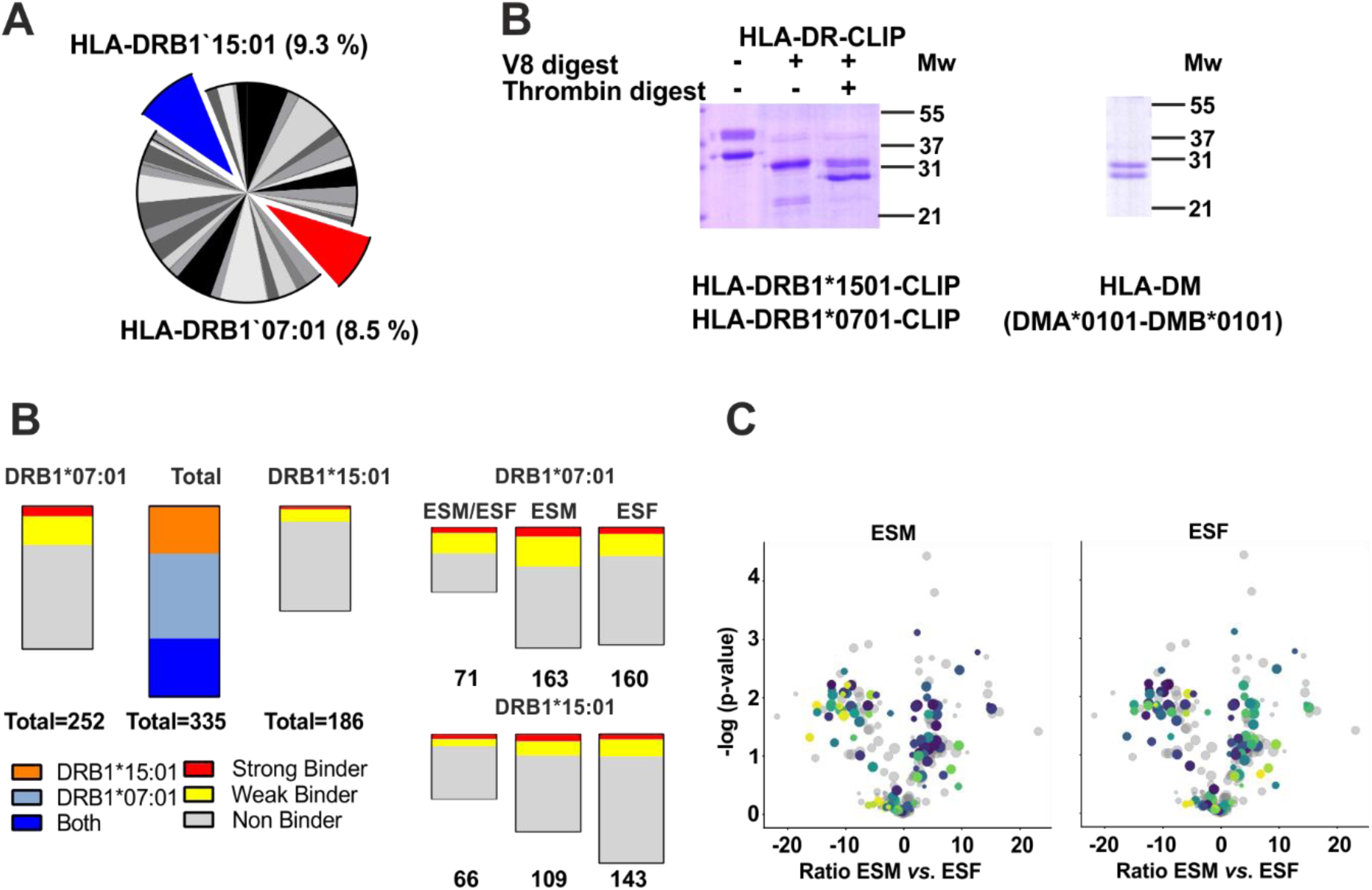
Additional experimental considerations for the reconstituted *in vitro* antigen processing system. **A.** Pie chart showing the abundance of HLA-DRB1 allotypes in the 1000 Genomes project. Each allotype is shown as a different color and full details of full are described in Gourraud, P.-A. et al. (2014). The two most abundant allotypes are DRB1*07:01 and DRB1*15:01. **B.** Details on the expression and purification of recombinant MHCII molecules and their processing for their use. **C.** Detailed analysis of the epitopes found for each MHCII allotype and each antigenic source (ESF and ESM).Potential S.B. and W.B. are indicated. **D.** Mapping of the identified potential epitopes to their corresponding protein sources. Colored dots (according to legend in main Fig 1E) indicate the protein sources that yield experimental epitopes

**Fig. S2.**
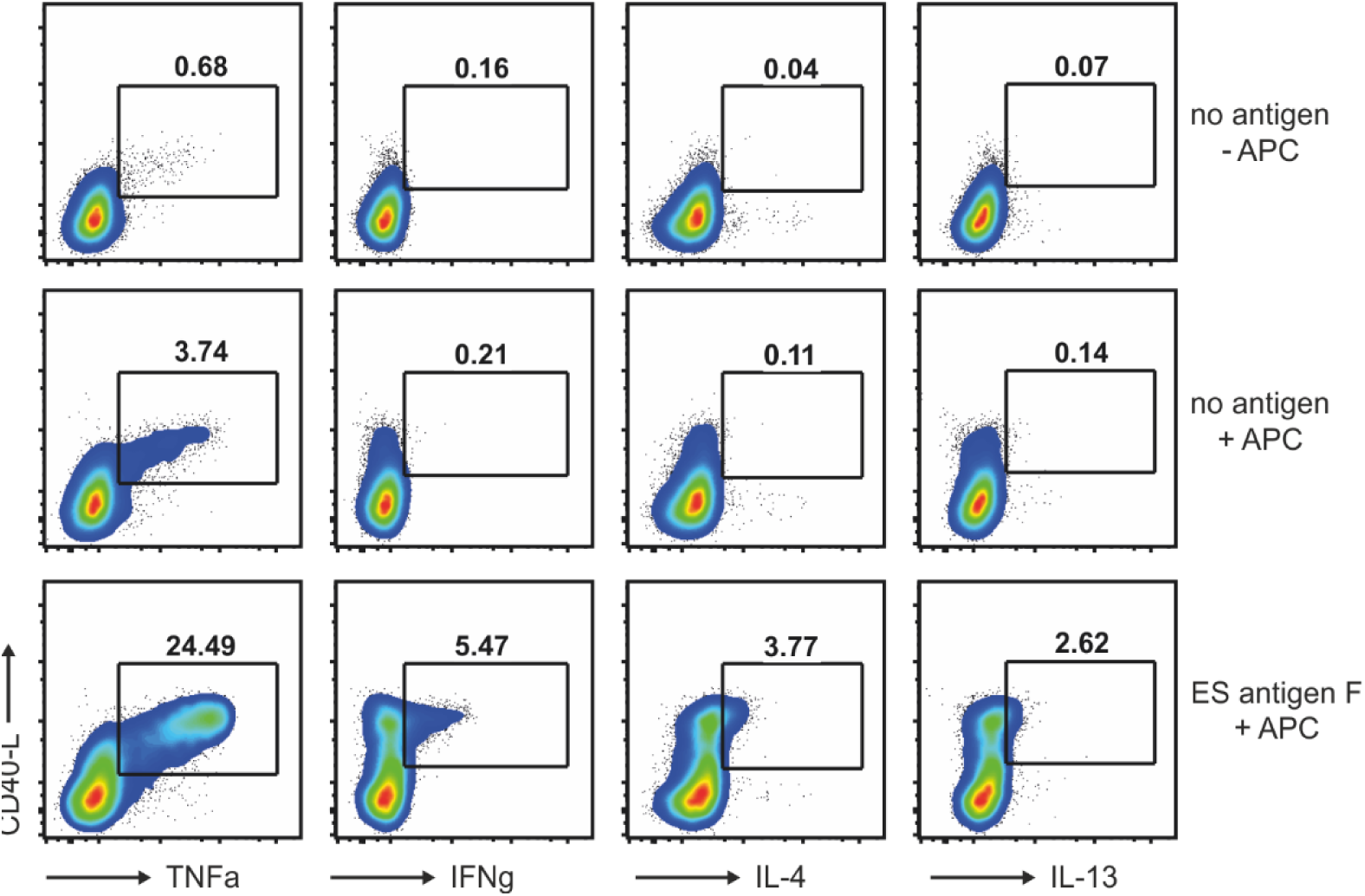
*Ascaris* ESF-specific T cell line following ESF restimulation analyzed for CD-40L/ cytokine co-expression and compared to no antigen and no antigen/no APC controls.

